# Cell type distribution of intrathecal antisense oligonucleotide activity in deep brain regions of non-human primates

**DOI:** 10.1101/2025.05.20.655163

**Authors:** Jeannine A Frei, Juliana E Gentile, Yuan Lian, Meredith A Mortberg, Juliana Capitanio, Paymaan Jafar-nejad, Sonia M Vallabh, Hien T Zhao, Eric Vallabh Minikel

## Abstract

Intrathecally administered RNase H1-active gapmer antisense oligonucleotides (ASOs) are promising therapeutics for brain diseases where lowering the expression of one target gene is expected to be therapeutically beneficial. Such ASOs are active, to varying degrees, across most or all cell types in the cortex and cerebellum of mouse and non-human primate (NHP), brain regions with substantial drug accumulation. Intrathecally delivered ASOs, however, exhibit a gradient of exposure across the brain, with more limited drug accumulation and weaker target engagement in deep brain regions of NHP. Here, we profiled the activity of a tool ASO against *Malat1* in three deep brain regions of NHP: thalamus, caudate, and putamen. All neuronal subtypes exhibited knockdown similar to, or deeper than, the bulk tissue. Among non-neuronal cells, knockdown was deepest in microglia and weakest in endothelial stalk. Overall, we observed broad target engagement across all cell types detected, supporting the relevance of intrathecal ASOs to diseases with deep brain involvement.

## Introduction

Lowering the expression of a specific target gene in the central nervous system (CNS) is expected to be therapeutically beneficial across a range of diseases^1^. Antisense oligonucleotides (ASOs) with gapmer design, where central region is unmodified DNA with 5’ and 3’ wing residues containing 2’ sugar modifications, can trigger RNase H1 degradation of a target RNA, lowering its expression^1^. More than 10 such gapmer ASOs administered intrathecally have entered clinical development for brain diseases, and one — tofersen, targeting *SOD1* — has been approved.

Available autopsy data from humans who received intrathecally administered ASOs support relatively broad brain distribution^2,3^, but as such data remain limited, our understanding of intrathecal ASO pharmacology comes primarily from pharmacology and toxicology studies in non-human primates (NHP) such as cynomolgus macaques, whose brain is ∼5% the mass of a human brain, and ∼200 times larger than a mouse brain^4^. An intrathecal ASO against *Malat1* distributed broadly throughout the brain of cynomolgus macaques, albeit with drug accumulation more than an order of magnitude lower in deep brain regions than many areas of the cortex and spinal cord^5^. Accordingly, target engagement in deep brain regions was also more modest. An intrathecal ASO against *PRNP* achieved broad knockdown across various cell types in the cynomolgus macaque cortex and cerebellum, with a pattern of activity similar to that observed in mouse^6^. Nevertheless, cortex and cerebellum exhibited strong drug accumulation in that study, with 21 and 16 µg/g of ASO respectively. The cell type specificity of gapmer ASO activity in deep brain regions with more limited drug concentration has not yet been evaluated. Here, we use single nucleus sequencing to analyze tissue from an NHP model, cynomolgus macaques, treated intrathecally with a *Malat1* ASO in order to characterize target engagement across cell types in each brain region.

## Results

### Selection of brain regions for evaluation

We re-analyzed the previously reported^5^ pharmacokinetic (PK) and pharmacodynamic (PD) parameters of a *Malat1* ASO. In a dose-response study in mouse and rat, the median inhibitory concentration (IC_50_) in tissue varied by brain region from 0.1 to 4 µg/g (Figure S1). We examined the PK versus PD across brain regions in NHPs (Figure 1A) that each received 3 doses of 25 mg on study days 1, 14, and 28, and tissue was collected on day 42. While most CNS tissues exhibited <20% residual target RNA, 5 regions exhibited weaker target engagement (Figure 1A-B). The three regions of the basal ganglia — putamen, globus pallidus, and caudate — exhibited the lowest drug accumulation, with 3/4 animals having drug accumulation at the lower limit of quantification (LLQ) in caudate and in putamen (Figure 1C). Thalamus exhibited somewhat higher drug accumulation and deeper target engagement, intermediate between basal ganglia and cortex. Cerebellum was unique in having modest target engagement despite higher drug accumulation, likely due to the predominance of cerebellar granule neurons (CGNs) which exhibit comparatively weak ASO activity. Because cerebellum was already examined for a *PRNP* ASO^6^, we chose not to revisit it here. We selected thalamus, caudate, and putamen for further evaluation by single nucleus RNA sequencing (snRNA-seq).

**Figure 1.**
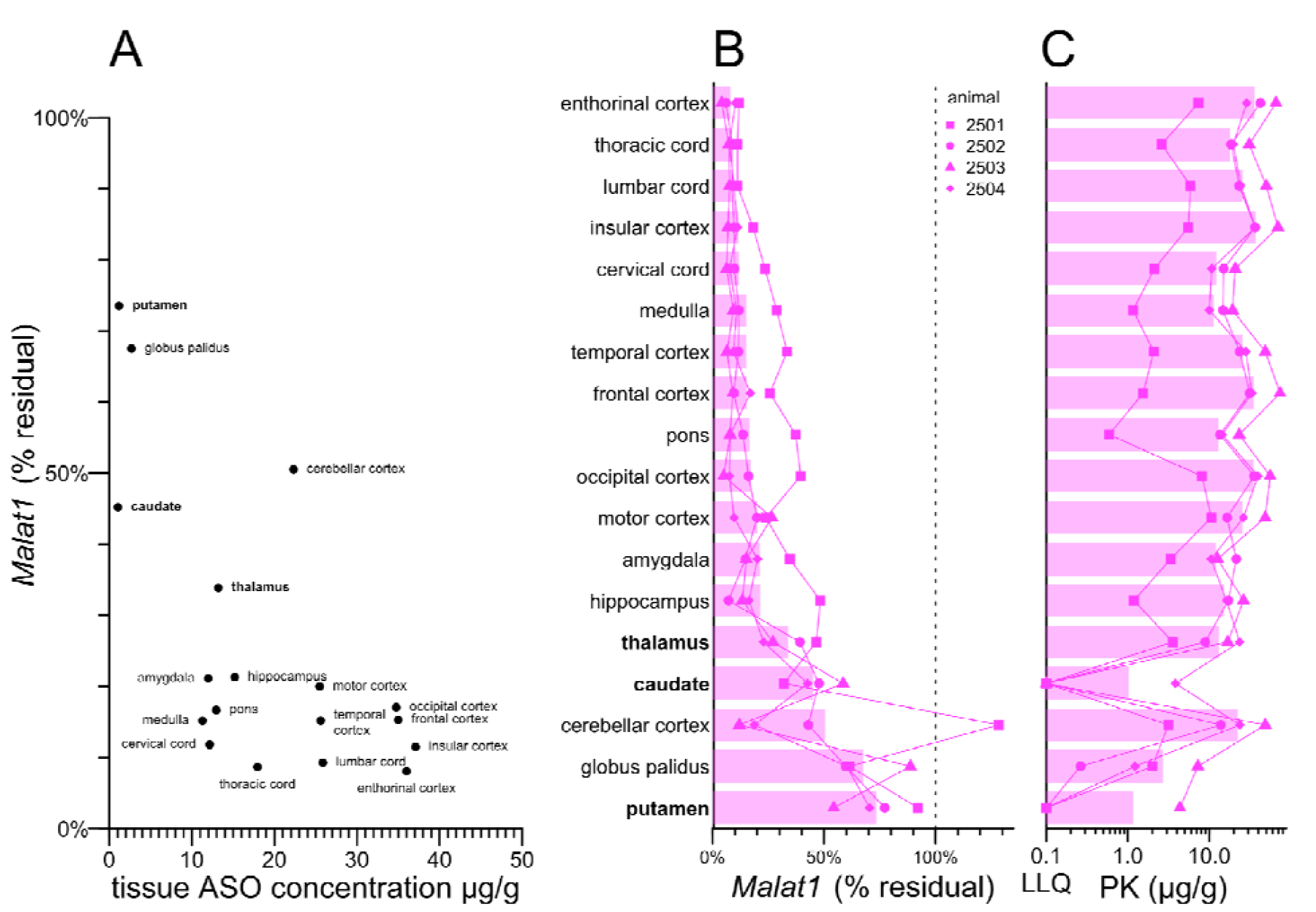
Pharmacokinetic and pharmacodynamic measurements across NHP CNS regions. All data in this figure are reproduced from Jafar-nejad & Powers et al. **A)** Mean pharmacokinetic measurements (tissue ASO concentration in µg/g) versus mean pharmacodynamic measurements (% residual Malat1 transcript by bulk qPCR). CNS regions selected for evaluation in this study are bolded. **B)** Pharmacodynamic measurements (% residual Malat1 transcript by bulk qPCR) for each ASO-treated animal in each region. **C)** Pharmacokinetic measurements (tissue ASO concentration in µg/g) for each ASO-treated animal in each region. In B-C animals are individual points, represented by different shapes for each animal; bars represent means.

Although all animals received the same dose of active drug, one animal (2501) exhibited more limited drug exposure, and weaker knockdown, across all CNS regions examined (Figure 1B-C) while another (2503) exhibited the highest drug exposure and deepest knockdown across most regions (Figure 1B-C). When the 4 animals were compared to one another across each of the CNS regions, higher drug exposure was significantly correlated with deeper target engagement (Figure S2).

### Identification of cell types in NHP deep brain

We performed snRNA-seq according to our previously published protocol^6^ on 3 brain regions each from 2 control NHP treated with aCSF and 4 NHP treated with *Malat1* ASO (18 samples total). We obtained transcriptomes from 130,553 single nuclei (Table S1). The 18 samples averaged 675 million reads each, mapping to 7,253 nuclei, with each nucleus having a median of 6,724 unique molecular identifiers (UMIs) nd 2,846 unique genes (Table S1). We used cell type-specific markers reported for mouse and marmoset^7^ to identify cell types (Figure 2 and S3). Across all three brain regions, we identified fast-spiking interneurons (IN) expressing parvalbumin (*PVALB*) and midbrain (MB)-derived IN expressing *OTX2, GATA3*, and *KIT*. In thalamus and caudate we also identified IN expressing somatostatin (*SST*). In thalamus, we found excitatory neurons expressing *RXFP1*, and caudate and putamen, *DRD1* and *DRD2*-positive spiny projection neurons (SPN), formerly known as medium spiny neurons. In addition to oligodendrocyte progenitor cells (OPC) and mature oligodendrocytes, in each brain region we also identified committed oligodendrocyte precursors / newly formed oligodendrocytes (COP/NFOL), typified by expression of *IFI44L* and a higher level of *ENPP6* expression than seen elsewhere. In caudate, we also identified radial glia by their particularly high expression of *VIM*.

**Figure 2.**
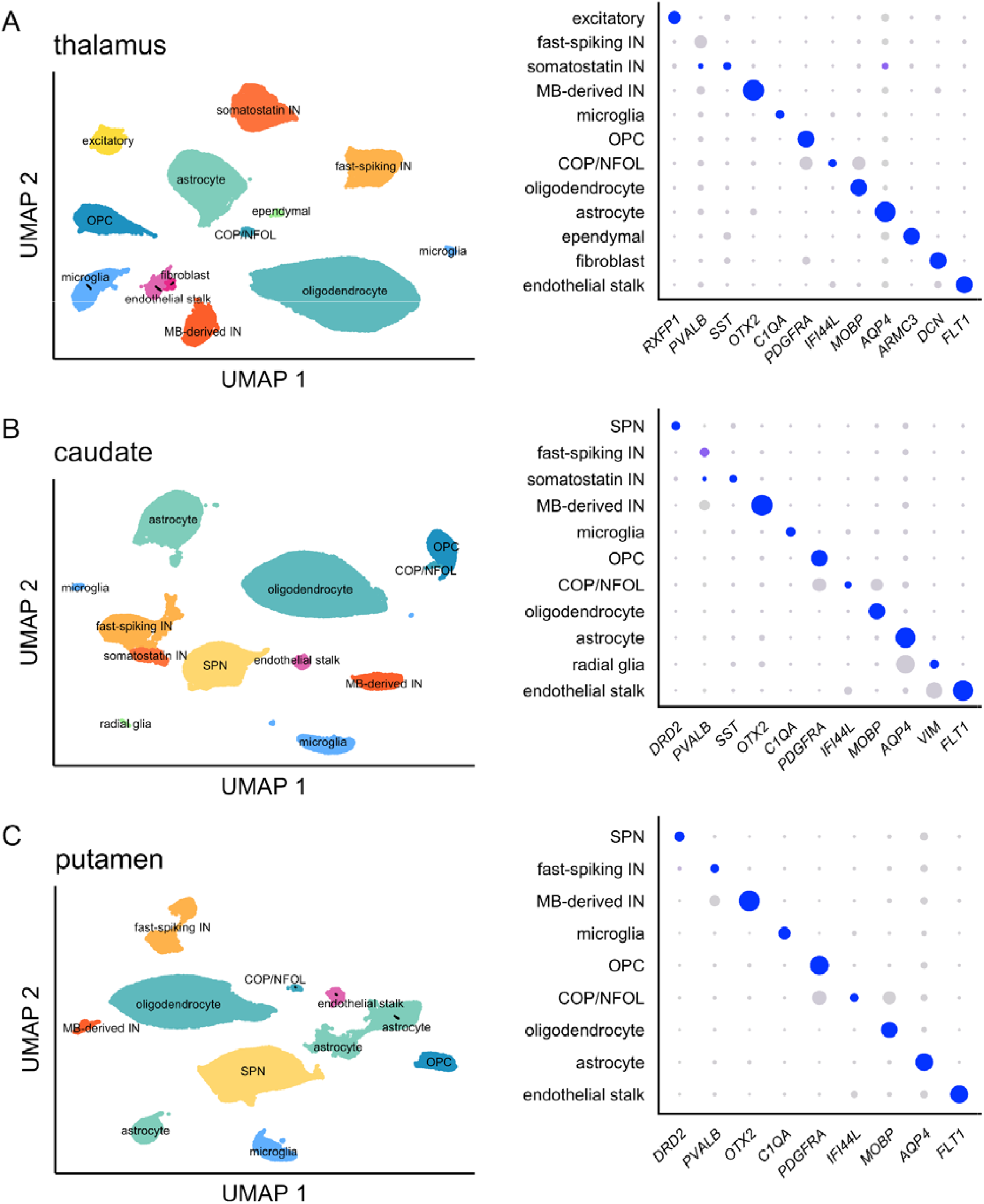
Cell types identified in NHP deep brain. Uniform manifold approximation and projection (UMAP) plots (left) and dotplots (right) for **A)** thalamus, **B)** caudate, and **C)** putamen. A larger panel of marker genes is included in expanded dotplots in Figure S3, and UMAPs for each individual animal are included in Figure S3.

While broadly distributed cell types such as oligodendrocytes and astrocytes were seen in roughly equal proportion across all animals, more regionally specific cell types were minimal or absent from some animals (Figure S4). For instance, in the thalamus, excitatory neurons were identified almost exclusively in 1/6 animals. Few or no SPNs were recovered in 2/6 animals in the caudate and 1/6 in the putamen. Dissection differences between animals likely account for these differences, as well as for the imperfect agreement between overall knockdown in single cell sequencing (Figure 3). For modeling the activity of the *Malat1* ASO, we only included animal + brain region + cell type combinations with ≥10 cells observed, and we only fit models for brain region + cell type combinations where at least 1 control and 2 treated animals met this criterion.

**Figure 3.**
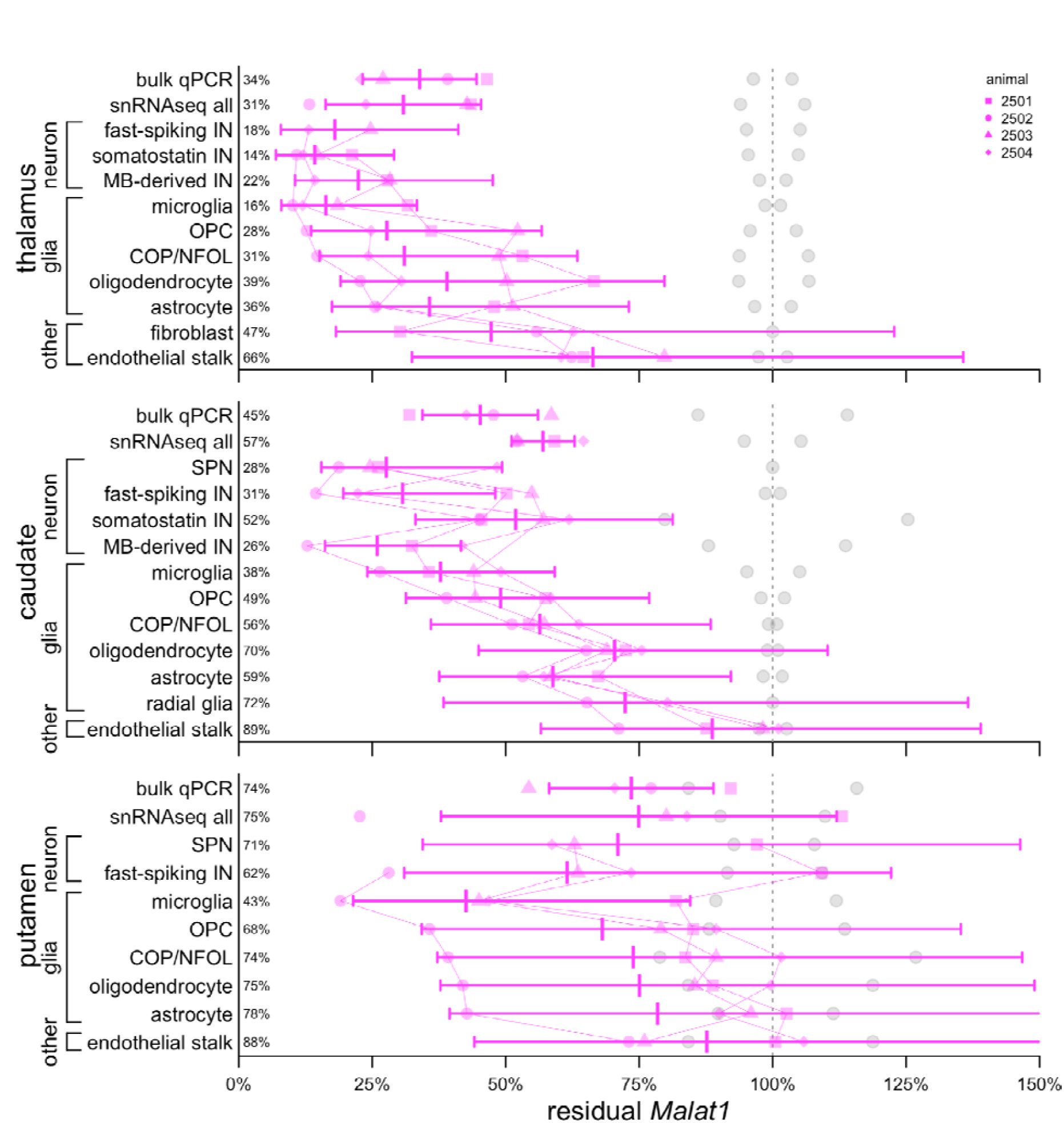
Target engagement by cell type in each NHP deep brain region. Gray points represent control animals (N=2), magenta points represent ASO-treated animals (N=4). Each animal’s data are represented by a unique shape and connected by thin lines between cell types. Bars represent means and 95% confidence intervals. Mean residual Malat1 is shown as a percentage at left.

Residual *Malat1* in bulk tissue averaged 31% in thalamus, 57% in caudate, and 75% in putamen by snRNA-seq, and 34%, 45%, and 75% respectively by qPCR (Figure 3). Just as observed previously in cortex and cerebellum^6^, all neuronal subtypes identified here exhibited residual target RNA at least as low as the bulk tissue, with putamen SPNs (71% residual) being nearest to the bulk (75%), while some neuronal subtypes were far lower than bulk, for instance, 14% and 18% residual for somatostatin and fast-spiking IN in thalamus, versus 31% in bulk tissue. Also consistent with previous results^6^, microglia exhibited relatively strong knockdown, deeper than any other glial cells, and in every brain region, endothelial stalk exhibited the weakest knockdown of any cell type. Overall, results were consistent with at least some knockdown in every detectable cell type in every brain region (Figure 3).

Histograms of *Malat1* UMIs for each region and cell type generally indicated that the entire distribution shifted left and remained unimodal (Figure S5). Only a few cell types in the putamen exhibited bimodal distributions of *Malat1* UMIs after ASO treatment, but this appeared to be due to different levels of knockdown in different animals. Distributions were consistent with knockdown in every individual cell, again as expected based on prior work^6^.

## Discussion

We previously reported a single nucleus characterization of ASO activity in the mouse brain as well as in the cortex and cerebellum of NHP^6^. Here we extend these results by examining three NHP deep brain regions in animals that received an intrathecal ASO against *Malat1*. Despite more limited drug exposure in these regions and weaker bulk knockdown, all our key results closely mirror the previous findings in cortex and cerebellum^6^: we observed ASO target engagement in every cell type detected; all neuronal subtypes have target engagement at least as deep as the bulk tissue; microglia exhibit especially deep knockdown while endothelial stalk cells exhibit the weakest knockdown; and distributions of target RNA counts across individual nuclei are consistent with ASO activity in every single cell. This last property of ASOs is contrast to a viral vectored zinc finger repressor, for which we observed nearly complete elimination of target transcript, only in transduced cells^8^. Given the number of ASO molecules administered in a dose, it is expected that every cell should exhibit some activity^6^.

Our study has several limitations. We examined just 18 samples from 6 animals treated with 1 ASO. The animals in this study received 3 doses of 25 mg ASO at 2-week intervals, which likely corresponds to greater drug exposure than achieved in clinical dosing regimens. The cynomolgus macaque brain is just 5% the mass of a human brain. We were unable to quantify knockdown in certain cell types because too few such cells were observed in some animals, likely due to dissection differences; the brain tissue from these animals has already been used for multiple prior studies, leaving imperfectly matched tissue punches for this work.

The finding of broad target engagement across cell types even in deep brain regions with more limited drug exposure may support the relevance of intrathecal dosing to treatment of whole brain diseases or diseases with particularly prominent deep brain involvement. Ultimately, the hypothesis that intrathecal ASO delivery can adequately address such diseases in the larger human brain can only be tested clinically.

## Methods

### ASO treatment of NHP

The *Malat1* ASO used here has been described previously^5,6^. Its sequence is <u>GComCoAoG</u>oGmCTGGTTATG<u>AomCoTmCA</u>, where o indicates phosphodiester linkages (other linkages are phosphorothioate), mC indicates 5-methylcytosine, and underline indicates 2’methoxylethyl. The animals examined here have been described previously^5^. Briefly, all were adult female *Macaca fascicularis* dosed intrathecally targeting the L4/L5 space with 25 mg ASO or artificial CSF (aCSF) delivered in a 1 mL volume over 1 minute, at days 1, 14, and 28 and harvested at day 42, at Charles River Laboratories Montreal under approval by the CRL Institutional Animal Care and Use Committee. 4 mm frozen coronal sections were punched to yield 2 mm diameter cylinders of tissue, cryostat dissected into quarters, with one quarter used for sequencing. Pharmacokinetic and pharmacodynamic (bulk qPCR) data used here were reproduced from a previous report. In instances where multiple samples were examined from one brain region for the same animal these were averaged across the whole brain region for each animal (example: thalamus VA/VL, thalamus VPM/VPL, and thalamus MD were averaged to yield a single measurement for thalamus).

### Single nucleus sequencing

We utilized the same wet lab protocol^9^ and computational pipeline^10^ described previously^6^. Briefly: frozen tissue was triturated in Kollidon VA64, Triton X-100, bovine serum albumin, and RNase inhibitor, passed through a 26-gauge needle, washed and pelleted, passed through a cell strainer, and flow sorted for DAPI signal. After cell counting, a volume estimated to contain 17,000 nuclei was submitted to the Broad Institute’s Genomics Platform for 10X library construction (3’ V3.1 NextGEM with Dual Indexing) was performed according to manufacturer instructions and sequenced (100 cycles) on an Illumina Novaseq 6000 S2. Data were analyzed on Terra.bio using Cumulus Cell Ranger 7.2.0 (Dockstore workflow github.com/lilab-bcb/cumulus/Cellranger:2.5.0) with flags –include introns and – secondary). Because *MALAT1* is not annotated the Ensembl Macaca fascicularis 6.0 transcriptome, we created our own custom reference. We downloaded the Macaca_fascicularis_6.0 GTF file, used cellranger mkgtf to filter for protein_coding, lncRNA, and antisense transcripts, manually added lines for *MALAT1* on the minus (-) strand spanning positions chr14:8898261-8905725, and finally compiled with cellranger mkref 8.0.1. Count matrices were then aggregated using Cell Ranger 7.1.0 (aggr with the –normalize flag set to none) to yield one UMI count matrix per species and brain region. Statistical analyses and data visualization were conducted using custom scripts in R 4.2.0.

### Cell type assignment

Preliminary cell types were assigned using scType^11^, and final cell type determinations were made after manual curation in Loupe Browser. In order to identify cell types, we used reported markers from projects SCP2706 and SCP2719 in the Single Cell Portal^7^ as well as previous publications on mouse, human, and NHP deep brain regions^11–14^.

### Statistical modeling

To estimate knockdown in each cell type, we utilized the same negative binomial model described previously. The model was specified as glm.nb(target umi ∼ celltype + celltype:treatment +offset(log(total umi))). The coefficient for each cell type-treatment interaction term (in natural logarithm space) was then exponentiated to yield the mean estimate of the residual target RNA in that cell type, and the 95% confidence interval was that mean estimate ±1.96 of the model’s standard error. Individual animal point estimates were obtained by adding the model’s residuals to each coefficient before exponentiating.

## Data and source code availability

An analytical dataset and R source code sufficient to reproduce these analyses will be made available at https://github.com/ericminikel/nhp_deepbrain_aso and the full sequencing dataset will be posted to the Single Cell Portal at https://singlecell.broadinstitute.org/

## Acknowledgments

This work was funded by Ionis Pharmaceuticals. We thank Briana Nobel for technical assistance with monkey tissue collection.

## Supplementary Figures

**Figure S1.**
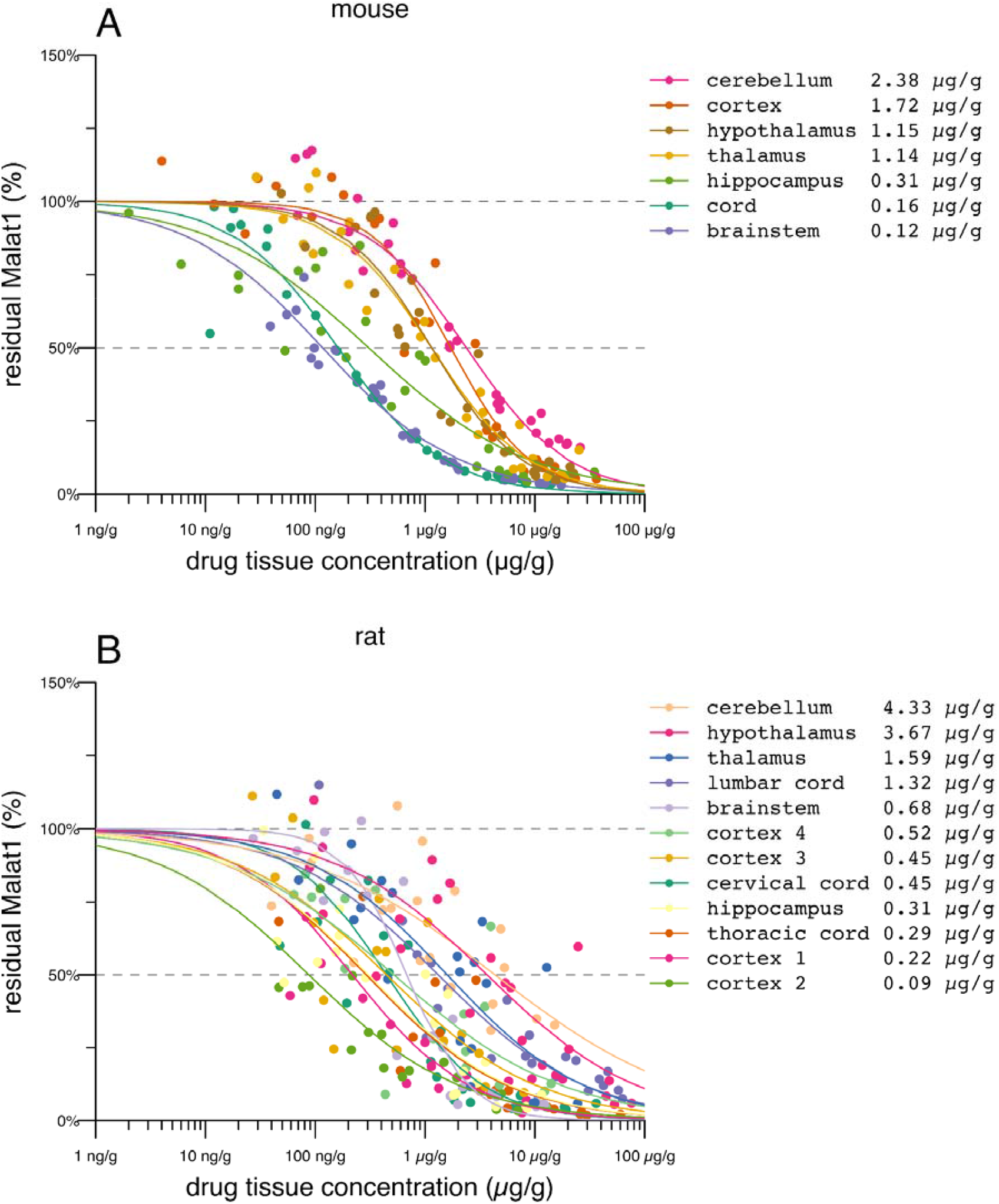
Pharmacokinetic/pharmacodynamic (PK/PD) relationships in mouse and rat for Malat1 ASO. Data are reproduced from Jafar-nejad & Powers et al for **A)** mouse and **B)** rat. For each brain region, a 4-parameter log-logistic dose-response curve is fit, with the response at zero dose fixed at 100% residual RNA and the response at infinite dose fixed at 0% residual RNA. Shown are dose-response curves and, in the legend, calculated IC_50_ values by tissue.

**Figure S2.**
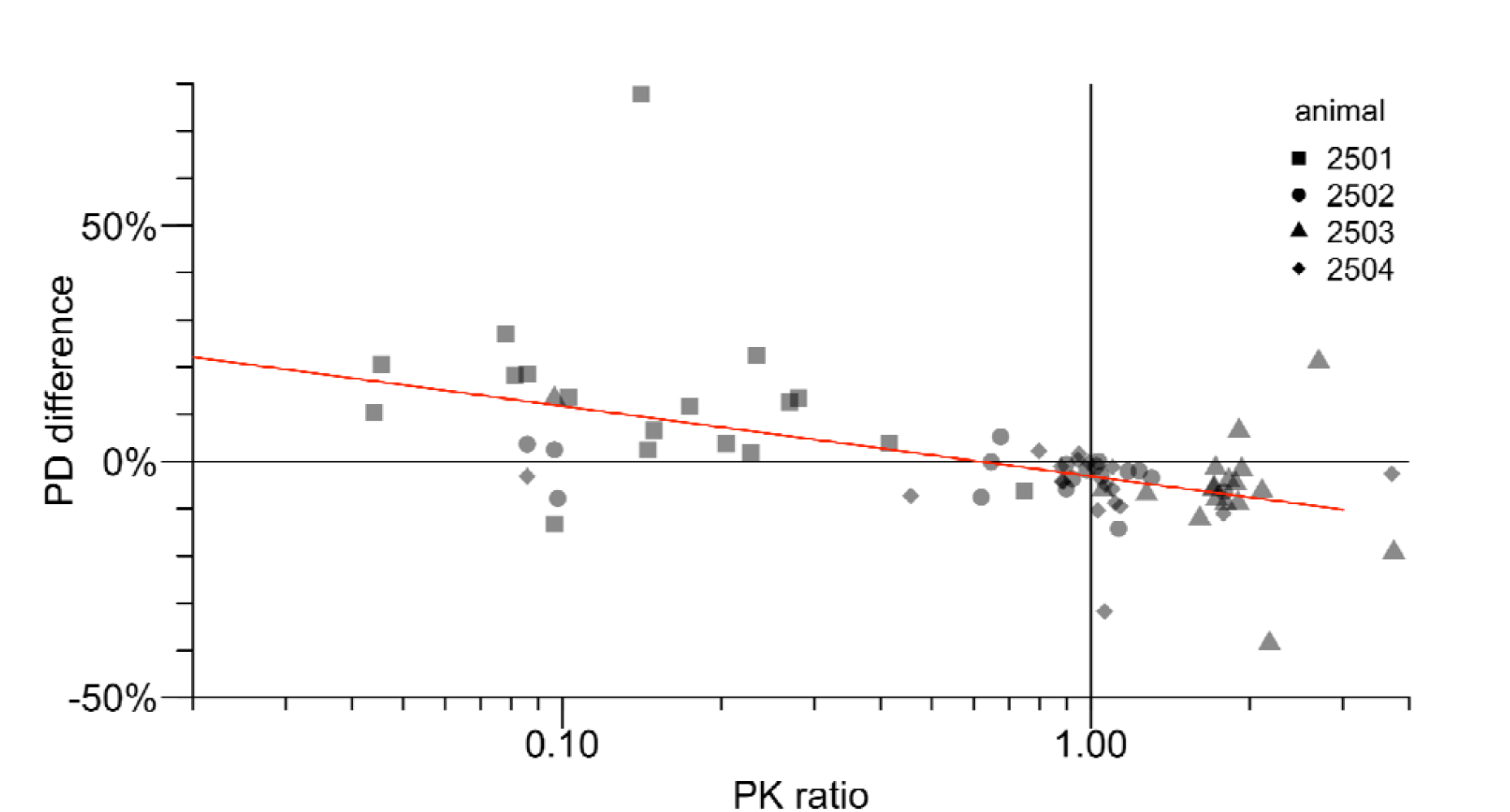
Correlation between individual animal pharmacokinetic and pharmacodynamic measurements. Each point represents one animal + CNS region combination, with shapes indicating which animal. The x axis is the ratio of the pharmacokinetic (PK) measurement in that animal (µg/g drug in tissue) to the average among the 4 animals — thus, 0.1 means the animal had 10% as much drug as the average animal. The y axis the difference in pharmacodynamic (PD) impact in that animal (% residual Malat1 transcript) in that animal versus the average among the 4 animals — thus, -10% means the animal had knockdown 10 percentage points deeper than the average knockdown, and +10% means the animal had knockdown 10% points less deep than the average knockdown. The red line is a log-linear model fit demonstrating a correlation whereby, across the individual animals and across CNS regions, higher drug concentration predicts deeper knockdown.

**FIgure S3.**
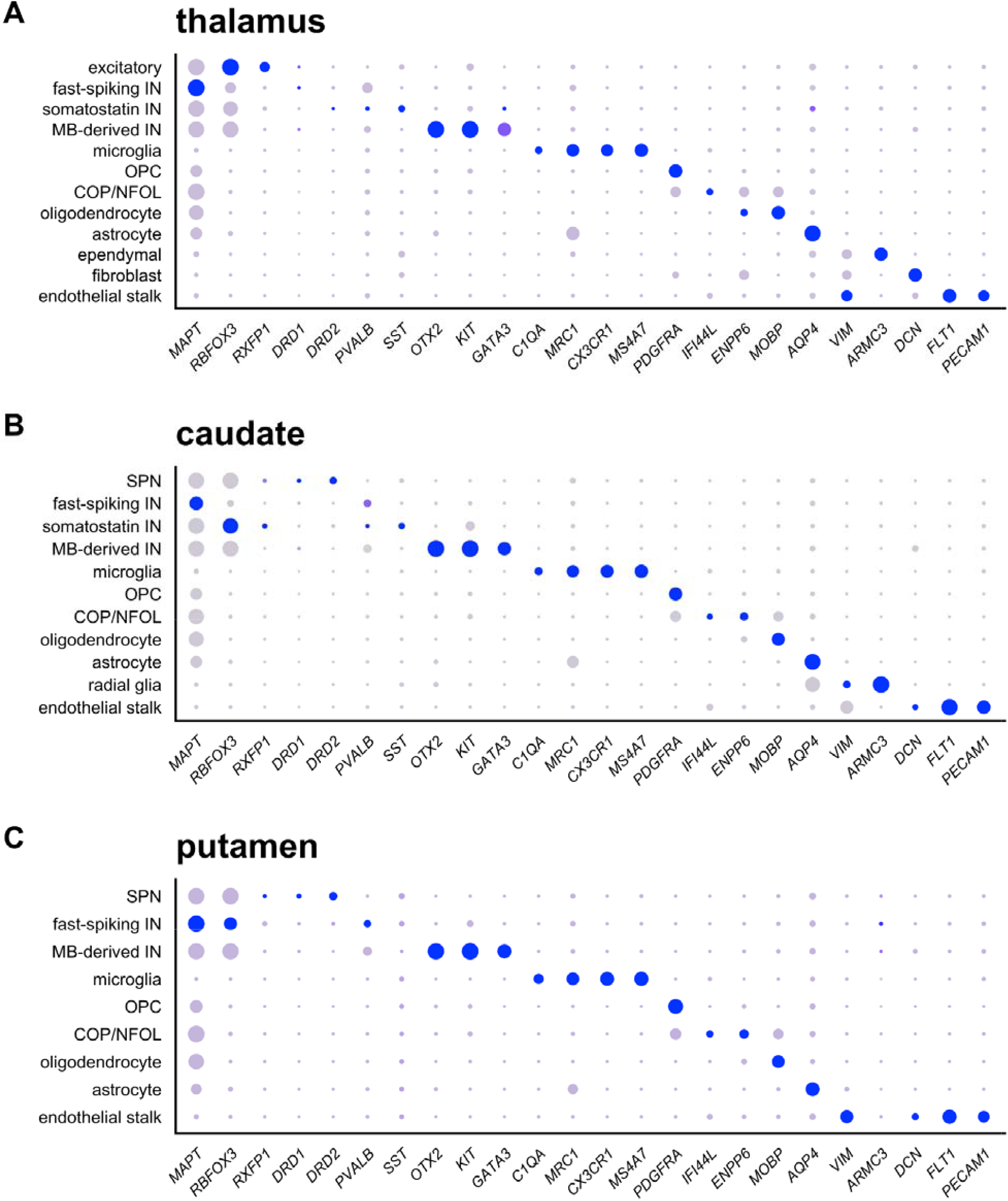
Dotplots with additional marker genes.

**Figure S4.**
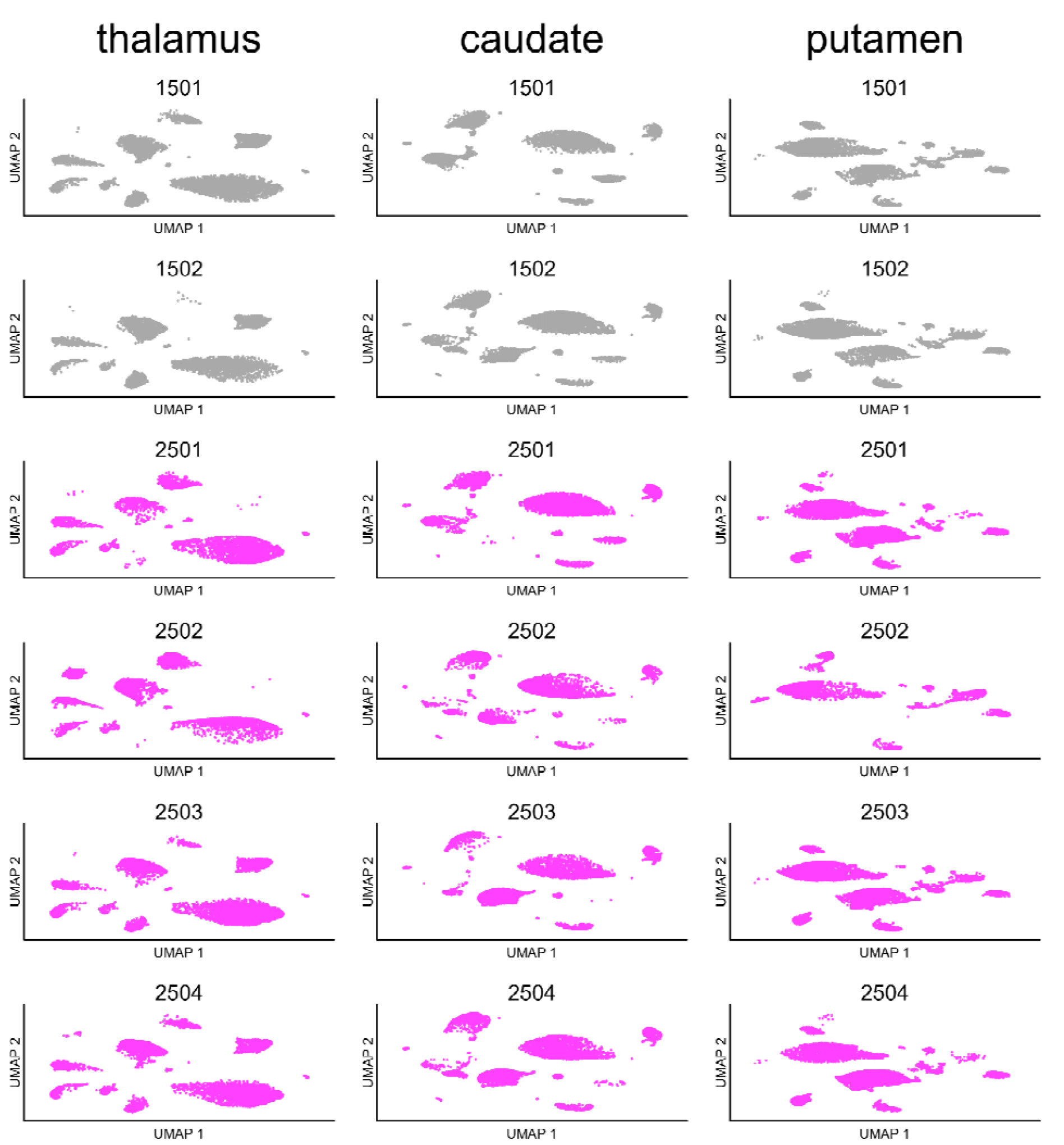
UMAP plots by individual animal. The absence of certain clusters of cells from certain animals corresponds to the dearth of certain cell types, perhaps due to regional dissection differences.

**Figure S5.**
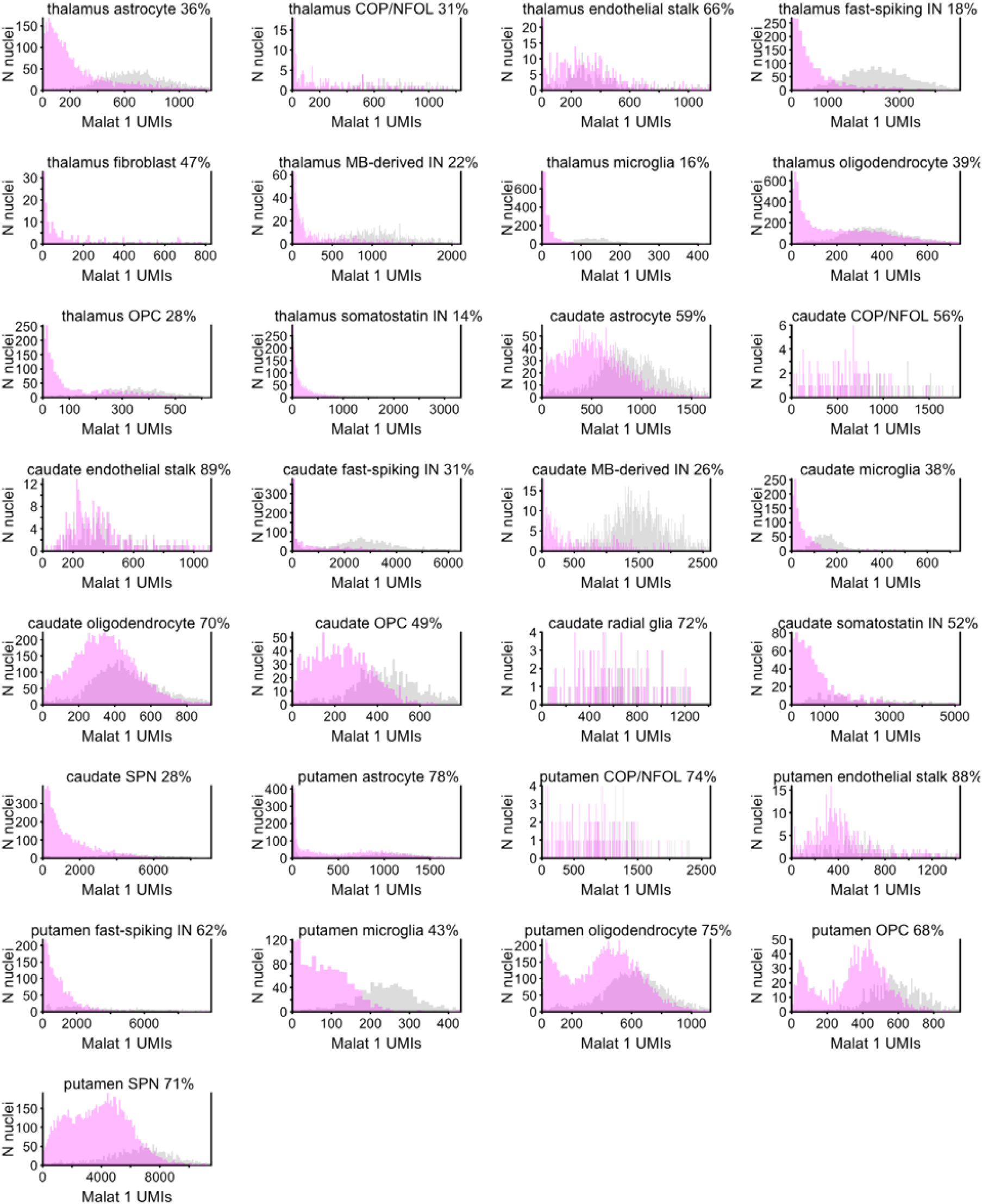
Histograms of Malat1 UMIs per individual nucleus by region and cell type. Each histogram represents 1 brain region and 1 cell type. Gray represents aCSF-treated animals while magenta represents ASO-treated animals. The x axis is the Malat1 UMIs per cell, and the y axis is the number of cells in that bin. Note that all distributions are roughly unimodal except for putamen oligodendrocyte and OPC. The left magenta peak in these distributions consists mostly of cells from animal 2502, which appeared to have relatively deeper knockdown than other animals across all cell types in putamen according to Figure 3.

